# A BASP1/BASP1-AS1 Axis Modulates Wnt and Notch Signaling to Balance Proliferation and Differentiation in Neuroblastoma Cells

**DOI:** 10.1101/2025.06.30.662295

**Authors:** Shubham Krishna, Mukta Kumari

## Abstract

Neuroblastoma exhibits significant intratumoral heterogeneity and resistance to differentiation therapy. We identify a regulatory axis between the protein-coding gene BASP1 and its antisense lncRNA BASP1-AS1 as a molecular switch between proliferation and neuronal differentiation in SH-SY5Y neuroblastoma cells. BASP1 maintains a proliferative, undifferentiated state by upregulating Wnt3a signaling and stemness-associated markers. Knockdown of BASP1 inhibits both proliferation and neuronal gene expression, implicating it as a context-specific oncogenic driver.

In contrast, BASP1-AS1 is transiently induced by retinoic acid (RA) and initiates early neuronal differentiation via DCX and MAP2 induction. BASP1-AS1 represses Wnt3a and activates Notch1, redirecting the signaling balance toward a differentiation-permissive state. A reciprocal suppression between BASP1 and BASP1-AS1 underlies a transition from Wnt3a to Wnt2 activity as differentiation progresses.

LiCl-mediated Wnt3a activation suppresses BASP1-AS1 and reinduces Sox2, highlighting Wnt3a’s role in maintaining stemness and therapy resistance. Post-RA BDNF treatment reinforces terminal differentiation, defined by high BASP1-AS1, DCX, and MAP2, and loss of proliferative signatures.

Together, these findings identify the BASP1/BASP1-AS1 axis as a central node integrating Wnt and Notch pathways to regulate plasticity and lineage progression in neuroblastoma. This axis represents a potential target for overcoming differentiation blockade and therapeutic resistance.

## Introduction

Neuroblastoma (NB) is the most common extracranial solid malignancy in children and accounts for approximately 15% of paediatric cancer-related deaths [1]. Originating from precursor cells of the sympathetic nervous system, these tumors frequently arise in the adrenal medulla and present with widespread metastasis at diagnosis [2]. Clinical management of NB relies heavily on risk stratification based on patient age, disease stage, chromosomal abnormalities, and MYCN gene amplification status [3]. High-risk NB, typically characterized by metastatic disease in children older than 18 months or tumors harbouring MYCN amplification, has a dismal prognosis, with five-year event-free survival rates below 50% [4]. The high relapse rate and severe long-term complications from current multimodal therapies highlight an urgent need for less toxic, mechanism-based therapeutic strategies [5].

Differentiation therapy has emerged as a promising approach to mitigate the malignancy of NB by promoting the terminal maturation of tumor cells. In contrast to conventional cytotoxic regimens, differentiation-based treatments hold the potential to improve survival outcomes while minimizing long-term toxicity [6]. Retinoic acid (RA), a key differentiation agent in current NB protocols, mediates its effects through nuclear retinoic acid receptors that bind specific DNA elements to regulate gene expression programs associated with neuronal lineage commitment [7–9]. Despite its clinical utility, the molecular mechanisms underpinning RA-induced differentiation remain incompletely defined, posing a barrier to the rational design of more effective differentiation therapies.

Wnt signaling, a central regulator of embryonic development, has also been implicated in NB pathogenesis. Dysregulation of the canonical Wnt/β-catenin pathway promotes proliferation, stemness, and chemoresistance in neuroblastoma cells [10–16], whereas activation of β-catenin independent Wnt signaling has been shown to suppress proliferation and support differentiation [17,18]. The mutually antagonistic interplay between β-catenin dependent and β-catenin independent Wnt pathways [19], along with cross-talk with additional signaling networks such as Notch [20], underscores the complex regulatory landscape governing NB cell fate. These insights point to the therapeutic potential of targeting Wnt signaling to modulate differentiation and overcome treatment resistance in high-risk NB [21].

To interrogate these mechanisms in vitro, the human neuroblastoma cell line SH-SY5Y is widely employed as a model of neuronal differentiation. These cells exhibit an undifferentiated, proliferative phenotype that can be reprogrammed into neuron-like cells under defined conditions, such as sequential treatment with RA followed by brain-derived neurotrophic factor (BDNF), leading to neurite outgrowth and expression of mature neuronal markers [22–24].

Neuronal differentiation in neuroblastoma is governed by intricate transcriptional programs and dynamic signaling transitions. Here, we demonstrate that the BASP1/BASP1-AS1 axis modulates biphasic Wnt and Notch signaling to control proliferation, lineage commitment, and terminal maturation in SH-SY5Y neuroblastoma cells. BASP1 promotes proliferation in undifferentiated cells, while BASP1-AS1 acts as a non-coding antagonist that facilitates Wnt2-mediated β-catenin-independent pathways and Notch1 activation in differentiating cells. Retinoic acid and BDNF sequentially induce transient transcriptional states, culminating in a mature, post-mitotic neuronal phenotype characterized by elevated DCX and MAP2, and repression of proliferation markers. Canonical Wnt activation by LiCl suppresses this axis and re-establishes stem-like features. Our findings define transcriptional thresholds that underpin differentiation and identify the BASP1/BASP1-AS1 circuit as a critical regulator of neuroblastoma cell fate. Previously, we had shown BASP1-AS1 as a critical lncRNA whose inhibition impairs neuroblastoma cancer stem cells (CSC) maintenance and proliferation. Neuroblastoma stem cells themselves are central to tumor aggressiveness, therapy resistance, and relapse, highlighting the importance of targeting regulators like BASP1-AS1 for improved treatment outcomes [25].

## Materials and methods

### Cell Culture and Differentiation

Undifferentiated SH-SY5Y human neuroblastoma cells were cultured in Dulbecco’s Modified Eagle Medium F12 (DMEM F12; Thermo Fisher Scientific) supplemented with 10% foetal bovine serum (FBS; Gibco). For neuronal differentiation, cells were treated with 10□μM retinoic acid (RA; Sigma-Aldrich) in complete DMEM, with media changes every other day. On fifth day of RA treatment, cells were washed with serum-free DMEM and subsequently maintained in serum-free DMEM supplemented with 50□ng/mL brain-derived neurotrophic factor (BDNF; PeproTech) for an additional eight days to promote neuronal maturation. Where indicated, recombinant DKK1 (Sigma-Aldrich) was added at a final concentration of 0.5□μg/mL. SH-SY5Y cells were seeded into appropriate culture plates and allowed to adhere for 24 hours. Following this incubation period, cells in the treatment group were exposed to 20 mM lithium chloride (LiCl), while control group cells were treated with an equivalent concentration (20 mM) of sodium chloride (NaCl). For subculturing and downstream applications, cells were gently dissociated using Accutase (Sigma-Aldrich).

### RNA Isolation and cDNA Synthesis

Total RNA was extracted using the standard Trizol–chloroform–isopropanol method. Briefly, cells were lysed in Trizol™ reagent (Invitrogen), and RNA was isolated through sequential addition of chloroform and isopropanol. Pelleted RNA was washed with 70% ethanol, air-dried, and resuspended in RNase-free water. RNA concentration and purity were assessed using a NanoDrop spectrophotometer, and only samples with an A260/280 ratio close to 2.0 were used. For cDNA synthesis, 1,000□ng of total RNA was reverse-transcribed using the High-Capacity cDNA Reverse Transcription Kit (Applied Biosystems) according to the manufacturer’s protocol. Reactions were performed at 25□°C for 10□min, 37□°C for 2□h, and 85□°C for 5□min, followed by a 4□°C hold. Synthesized cDNA was stored at −20□°C until further use.

### Quantitative Real-Time PCR (qRT-PCR)

qRT-PCR was conducted using SYBR™ Green chemistry (Applied Biosystems) on a Rotor-Gene Q platform (Qiagen). Each 20□μL reaction contained 10□μL SYBR Green Master Mix, 1□μL cDNA template, 0.5□μL each of forward and reverse primers (10□μM), and 8□μL nuclease-free water. Thermocycling parameters included initial denaturation at 95□°C for 5□min, followed by 40–45 cycles of 95□°C for 15□s, 62□°C for 30□s, and 72□°C for 1□min. Relative gene expression levels were calculated using the ΔΔCt method, normalized to GAPDH or PPIA. All experiments were conducted in at least three independent biological replicates. Primer sequences are listed in Supplementary Table 1.

### siRNA Transfection

For siRNA-mediated knockdown, SH-SY5Y cells were seeded in 6-well plates and transfected at 60–70% confluency with 80□pmol of siRNA targeting BASP1-AS1 or BASP1 (Catalogue No. HSS138797 or 4392421, Thermofisher) using Lipofectamine™ RNAiMAX or 3000 (Invitrogen), as per manufacturer’s instructions. A non-targeting scrambled siRNA (Invitrogen) was used as a negative control. For combined transfection and differentiation experiments, cells were transfected on Day 0 and four-hours post-transfection, the medium was replaced with differentiation-inducing medium containing 10□μM RA. Transfected cells were maintained under differentiation conditions for 72□h. Cells were also transfected during the differentiation process on day 1. Knockdown efficiency was confirmed at specified time points using qRT-PCR.

### Western Blotting

Cells were lysed in RIPA buffer supplemented with protease inhibitors (Roche). Total protein concentrations were quantified using the BCA Protein Assay Kit (Thermo Fisher Scientific). Equal amounts of protein (30□μg) were resolved by SDS-PAGE and transferred to nitrocellulose membranes (Millipore). Membranes were blocked with 5% non-fat milk in TBS-T (Tris-buffered saline with 0.1% Tween-20) for 1□h at room temperature and incubated overnight at 4□°C with β-catenin (Milipore-05-665), DCX (Santacruz-271390), Sox2 (Abcam-97959) primary antibodies. After washing, membranes were incubated with HRP-conjugated secondary antibodies (Vector Laboratories, Catalog No. PI-2000, PI-1000) for 1□h at room temperature. Protein bands were visualized using enhanced chemiluminescence (ECL; Milipore) and imaged using a ChemiDoc system. Densitometric quantification was performed with ImageJ software. Protein levels were normalized to α-tubulin as a loading control. All Western blot experiments were conducted in at least three independent biological replicates.

### Statistical Analysis

All quantitative data are presented as mean ± standard deviation (SD) from at least three independent biological replicates. Statistical comparisons between two groups were performed using Student’s t-test. Significance levels were denoted as follows: P < 0.05 (*), P < 0.005 (**), P < 0.0005 (***), and non-significant differences as # (p ≥ 0.05).

**Table 1.**
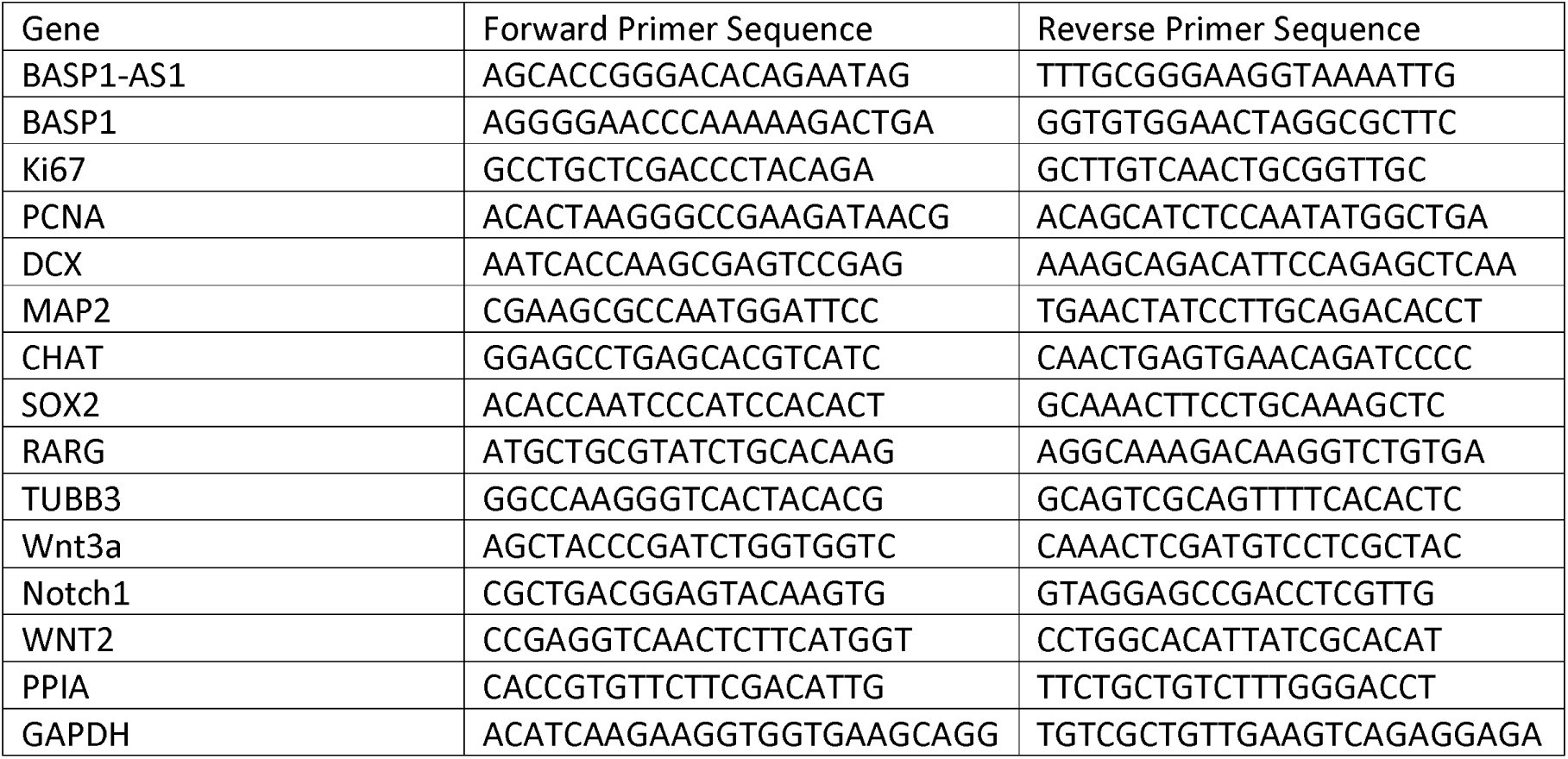
Primer Sequences.

## Results

### 1. BASP1 regulates SH-SY5Y cell morphology, proliferation, and differentiation through Wnt3a signaling

To investigate the role of BASP1 in undifferentiated SH-SY5Y cells, cells were transfected with BASP1-specific siRNA. Compared to cells treated with scrambled siRNA, BASP1 knockdown cells exhibited a more spread-out morphology and reduced cell density, indicative of decreased proliferative capacity. Cells treated with scrambled siRNA appeared denser, more compact, and rounded – hallmarks of undifferentiated, proliferative SH-SY5Y cells. (Figure 1A). Furthermore, BASP1 silencing led to a significant decrease in the expression of proliferation markers Ki67 and PCNA, as well as neuronal differentiation markers DCX and MAP2 (Figure 1B-F). Mechanistically, BASP1 depletion led to significant downregulation of Wnt3a, while activity of the Notch1 signaling pathway remained unaffected (Figure 1G-H). BASP1-AS1 depletion depleted the oncogenic Wnt3a dependent BASP1 expression (Figure 1I-J).

**Figure 1.**
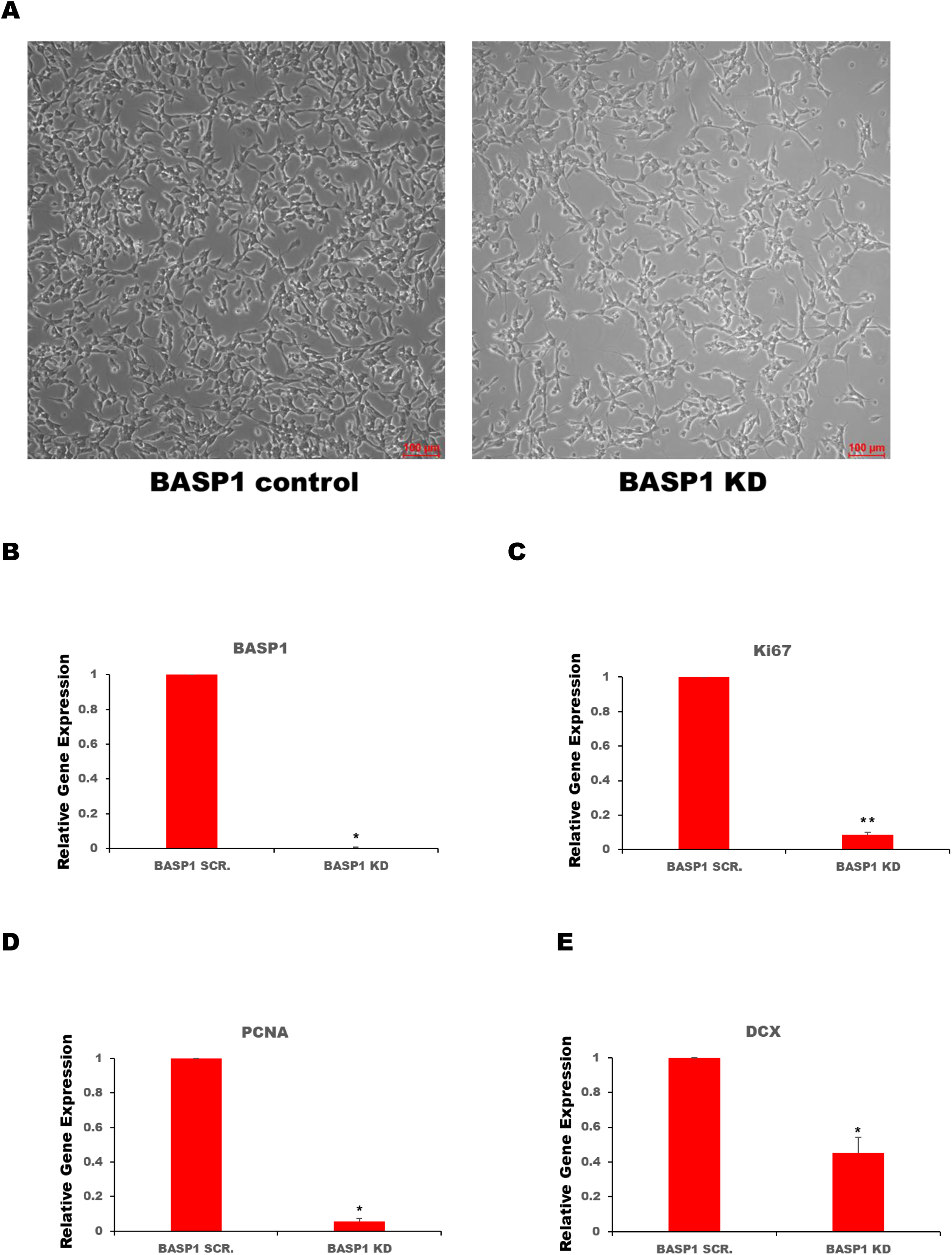
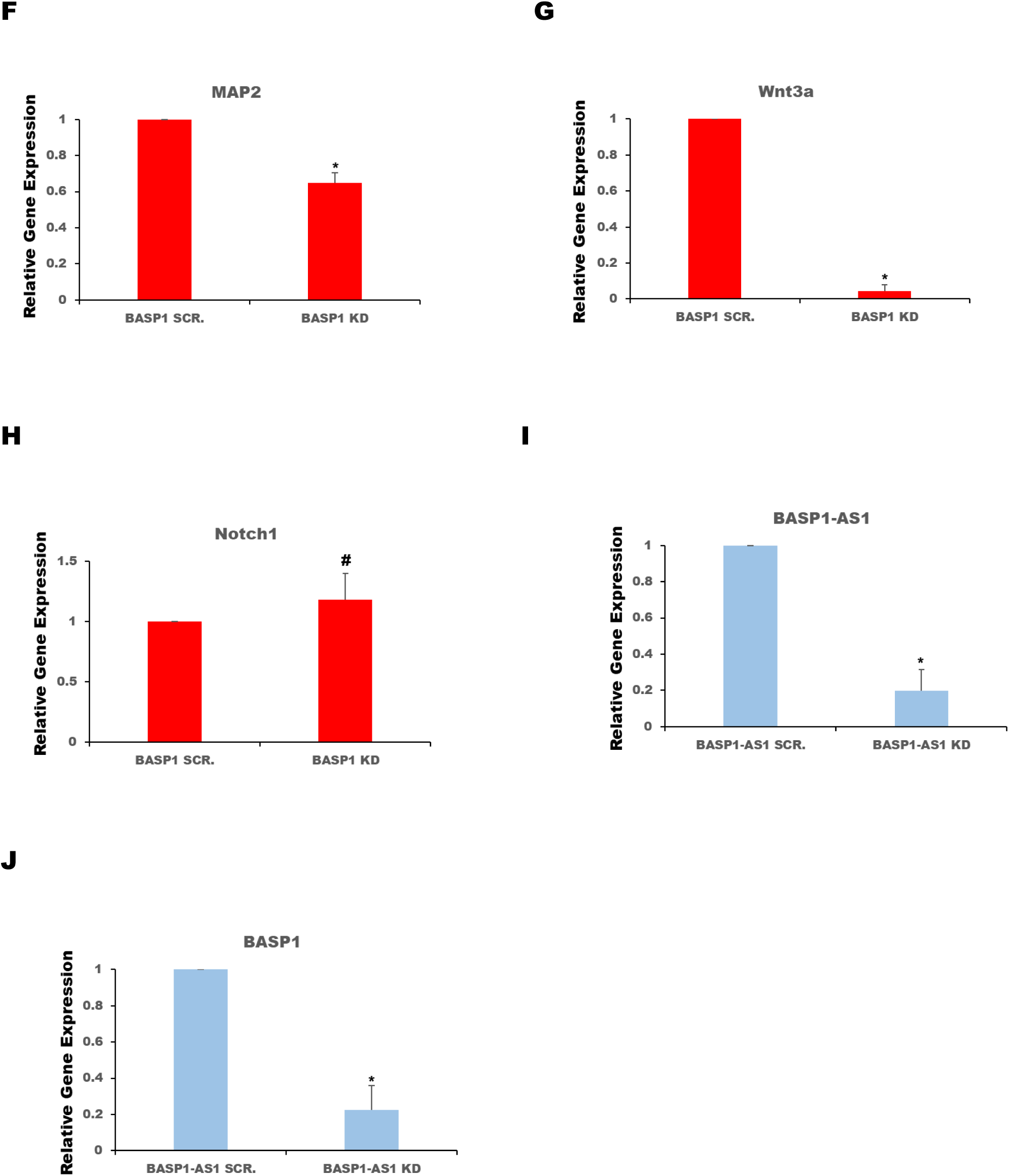
| BASP1 regulates SH-SY5Y cell morphology, proliferation, and differentiation through Wnt3a signaling. (A) Phase-contrast images of undifferentiated SH-SY5Y cells transfected with either scrambled siRNA (control) or BASP1-specific siRNA. BASP1 knockdown cells show a more spread-out morphology and reduced cell density compared to control cells, which remain rounded and compact. (B–D) Quantification of BASP1 (B) Ki67 (C) and PCNA (D) mRNA following BASP1 knockdown, indicating reduced proliferative capacity. (E–F) Expression levels of neuronal differentiation markers DCX (E) and MAP2 (F) are significantly decreased upon BASP1 silencing. (G–H) RT-qPCR of canonical Wnt3a (G) and Notch1 (H) signaling components. Wnt3a expression is significantly downregulated following BASP1 depletion, while Notch1 remains unchanged. (I–J) Silencing of BASP1-AS1 leads to concomitant downregulation of Wnt3a (I) and BASP1 (J) expression, supporting a Wnt3a-dependent regulation of BASP1. All data represent mean ± SD from at least three independent experiments. Statistical significance was determined by Student’s t-test. *P < 0.05, ** P < 0.005, ^#^ p ≥ 0.05

### 2. RA-Induced Differentiation in SH-SY5Y Cells follows Biphasic Activation-Resolution Dynamics

To assess the role of BASP1/BASP1-AS1 in neuroblastoma cell differentiation, SH-SY5Y cells were subjected to retinoic acid (RA)-induced differentiation over a 7-day period. As shown in Figure 2A, progressive neuronal differentiation was characterized by increased neurite length and complexity, alongside a transition of cell bodies from a rounded morphology to a more neuron-like, elongated shape. Over time, differentiated cells formed intricate neuritic networks indicative of advancing maturation.

**Figure 2.**
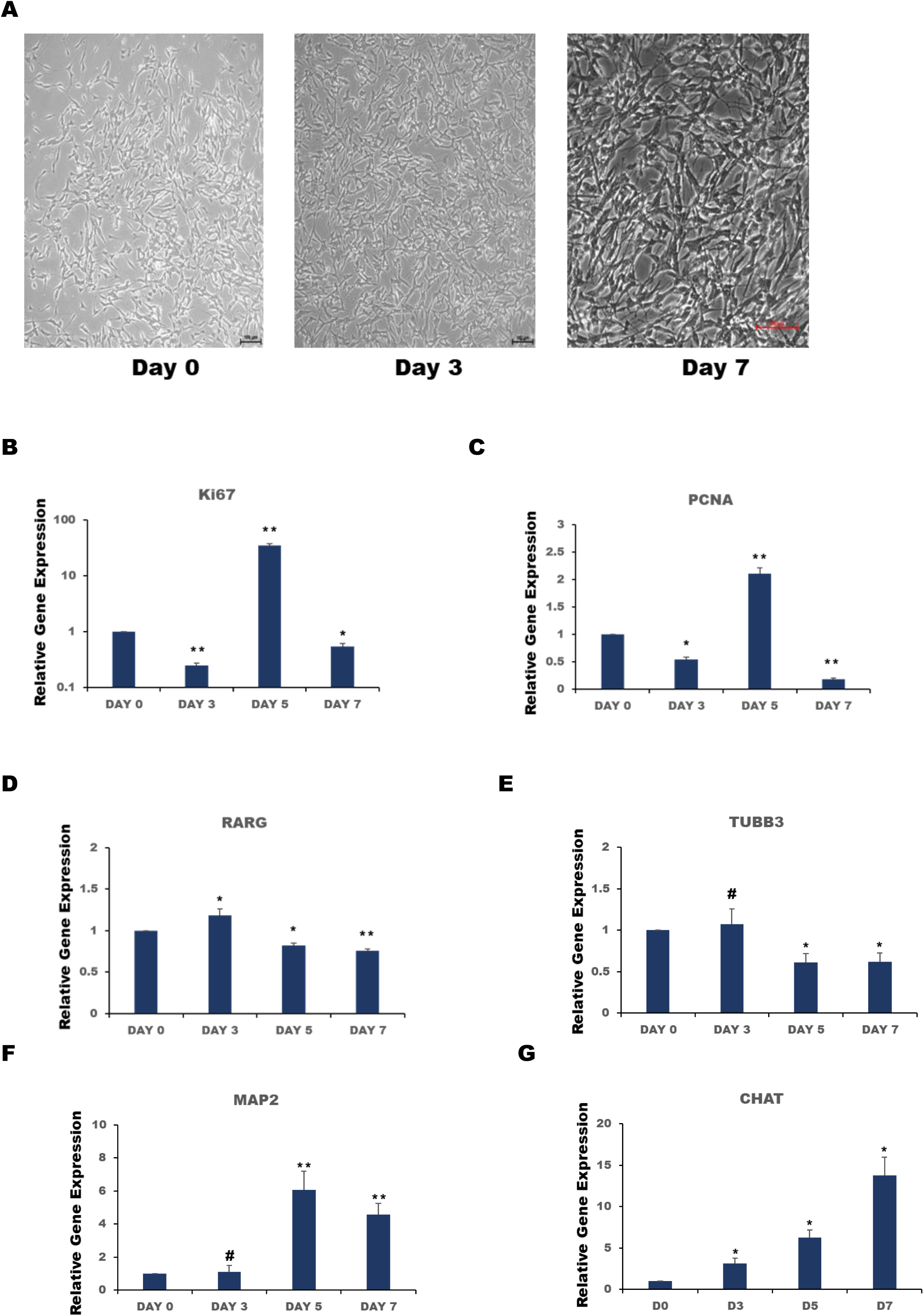
| RA-induced differentiation in SH-SY5Y cells follows biphasic activation-resolution dynamics. (A) Morphological changes observed during RA-induced differentiation of SH-SY5Y cells over a 7-day period. Differentiation is marked by increased neurite length, complexity, and a transition from rounded to elongated cell body morphology. Formation of interconnected neuritic networks is evident by day 7. (B–C) Quantitative analysis of proliferation markers Ki67 (B) and PCNA (C), both significantly downregulated by day 3, reflecting early cell cycle exit. (D) RARG expression peaks modestly (∼18% increase) at day 3, followed by a reduction below baseline by days 5 and 7, indicating a transient role in early neuronal lineage commitment. (E) βIII-tubulin (TUBB3) expression shows a ∼40% decrease by days 5 and 7, potentially reflecting cytoskeletal remodelling. (F–G) Terminal neuronal differentiation markers MAP2 (F) and CHAT (G) are significantly upregulated from day 5 onwards, consistent with acquisition of mature neuronal characteristics. All data represent mean ± SD from at least three independent experiments. Statistical significance was determined by Student’s t-test. *P < 0.05, ** P < 0.005, ^#^ p ≥ 0.05

Markers of cellular proliferation, including Ki67 and PCNA, were significantly reduced by day 3, indicating an early exit from the cell cycle. Ki67 levels dropped to ∼75% of undifferentiated levels (Figure 2B), with PCNA displaying a similar downregulation (Figure 2C), marking a transition toward a post-mitotic state – a defining feature of neuronal differentiation.

RARG, a key regulator of neuronal fate specification □26], showed a modest increase (∼18% over baseline) at day 3 (Figure 2D), followed by a steady decline below basal levels by days 5 and 7. This expression pattern suggests a temporally restricted role for RARG in early differentiation, possibly to initiate lineage commitment while subsequently repressing self-renewal programs.

The expression of βIII-tubulin (TUBB3), a neuron-specific cytoskeletal marker, showed a subtle, non-significant change on day 3 but declined by ∼40% by days 5 and 7 (Figure 2E). This reduction may contribute to cytoskeletal remodelling, promoting axonal and dendritic extension required for acquiring mature neuronal morphology [27,28].

Markers of terminal neuronal differentiation, including MAP2 and CHAT, were significantly upregulated by day 5 and remained elevated thereafter (Figure 2F-G), indicating progressive neuronal maturation and the initiation of functional synaptic properties.

These data support a model in which neurogenic cues operate within a biphasic framework that defines two key windows during neuroblastoma cell differentiation: an activation phase (Days 0–3) characterized by proliferative exit and initiation of neurogenic signaling, and a resolution phase (Days 4–7) marked by attenuation of early signals and the emergence of mature neuronal phenotypes.

### 3. Reciprocal regulation between BASP1 and BASP1-AS1 modulates Notch1 signaling and promotes neuronal marker expression

Protein expression analysis revealed that retinoic acid (RA) treatment led to an early progressive decline in β-catenin, DCX and Sox2 levels during differentiation, with these markers remaining consistently suppressed throughout the time course (Figure 3A-E).

**Figure 3.**
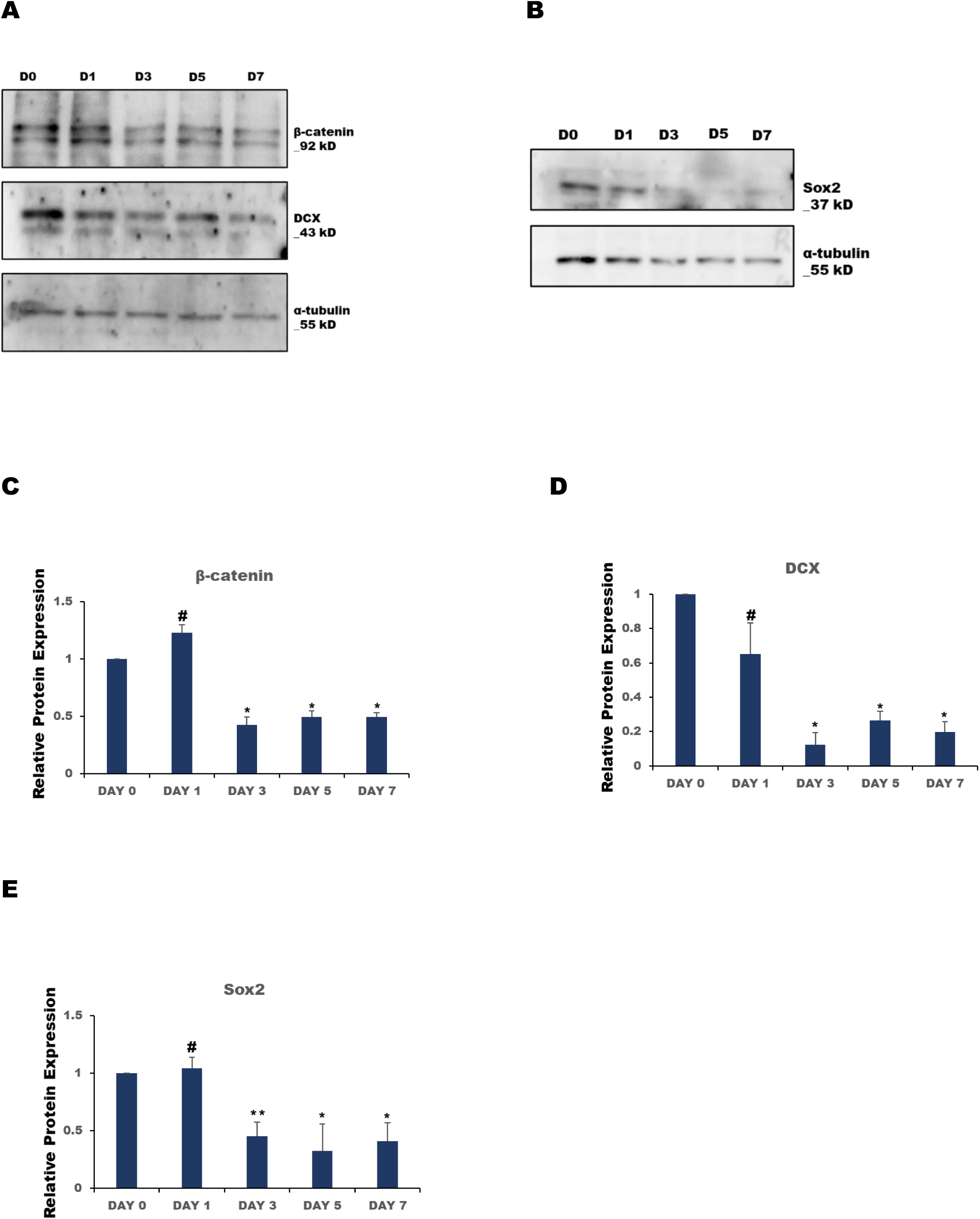
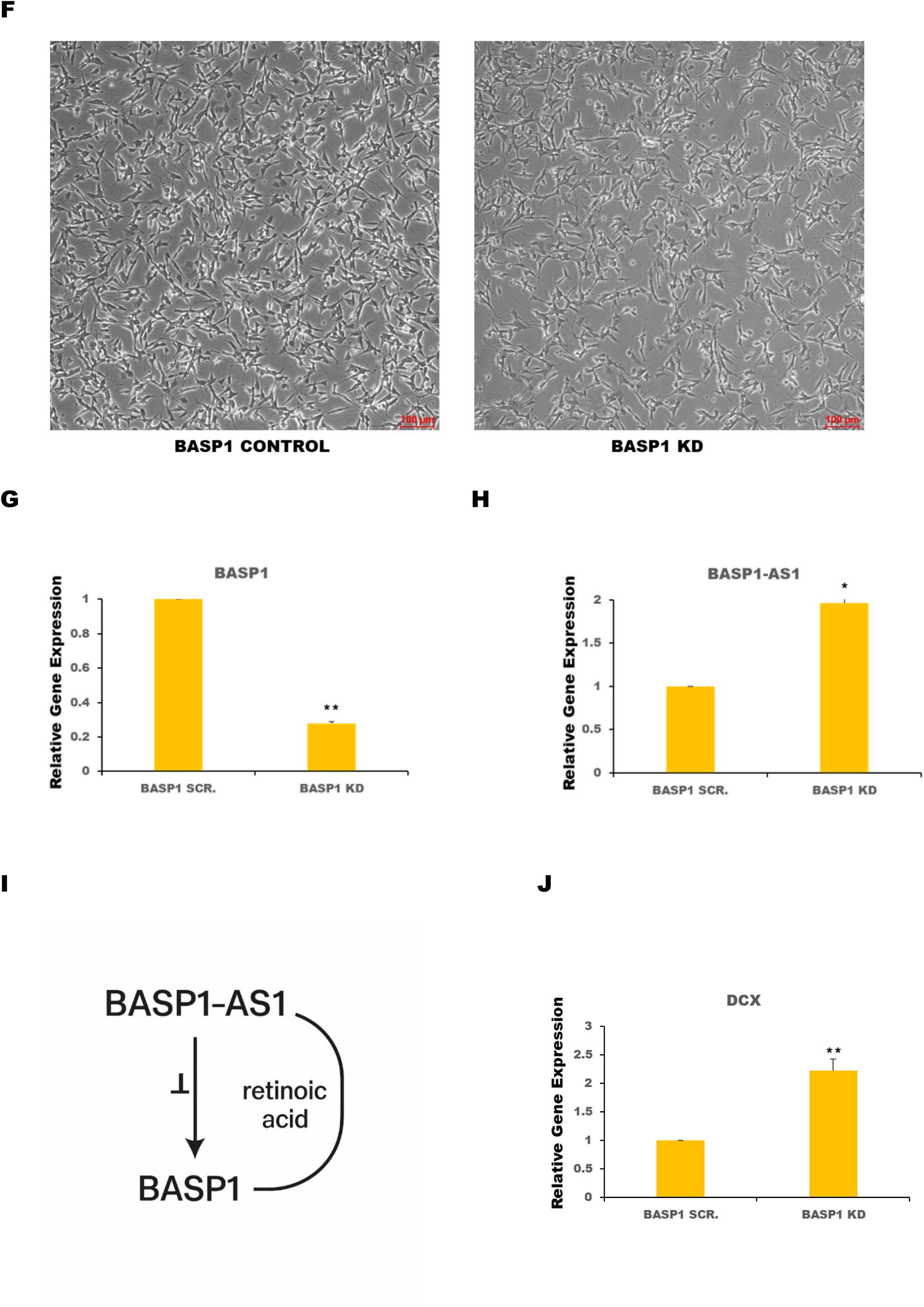
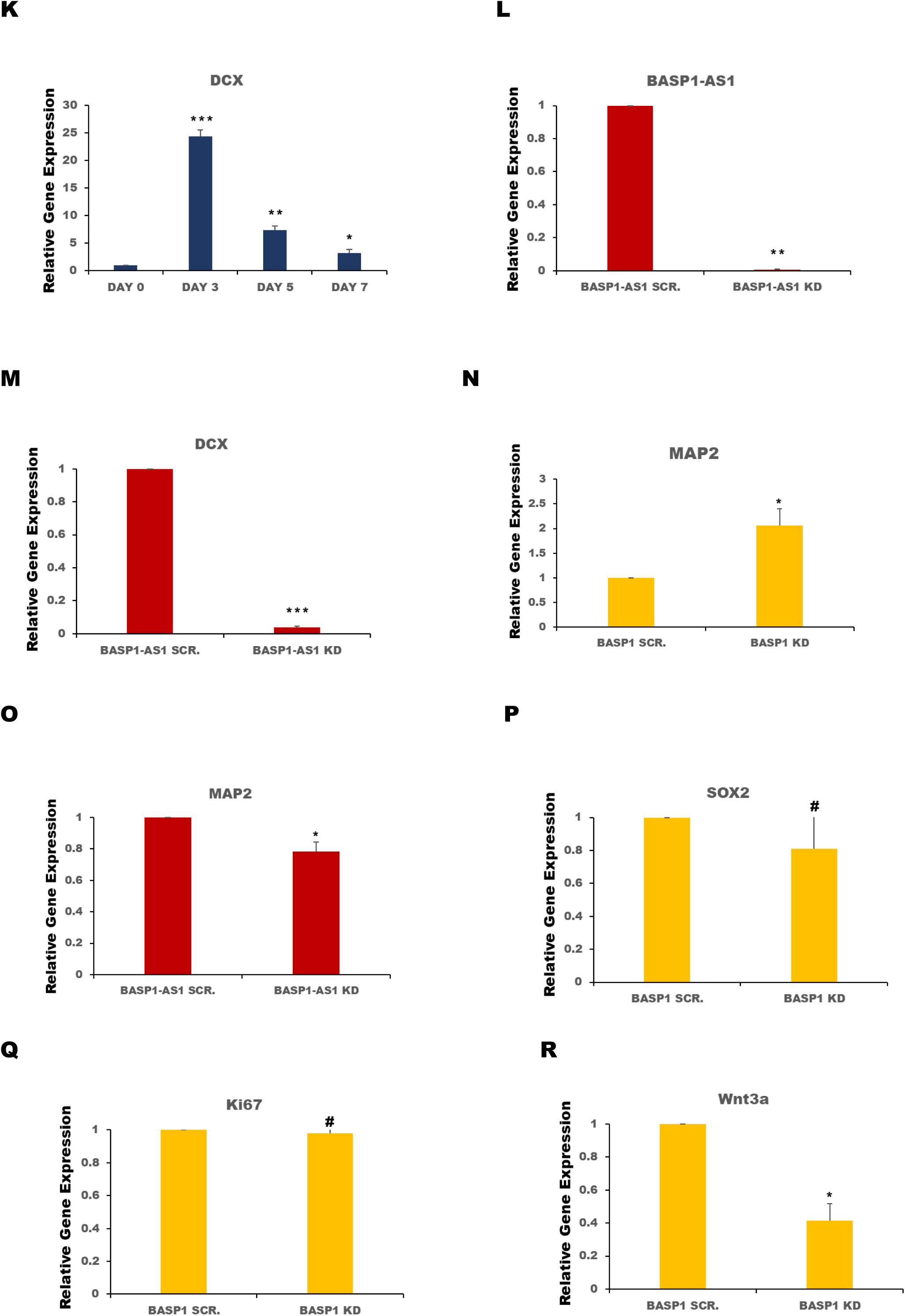
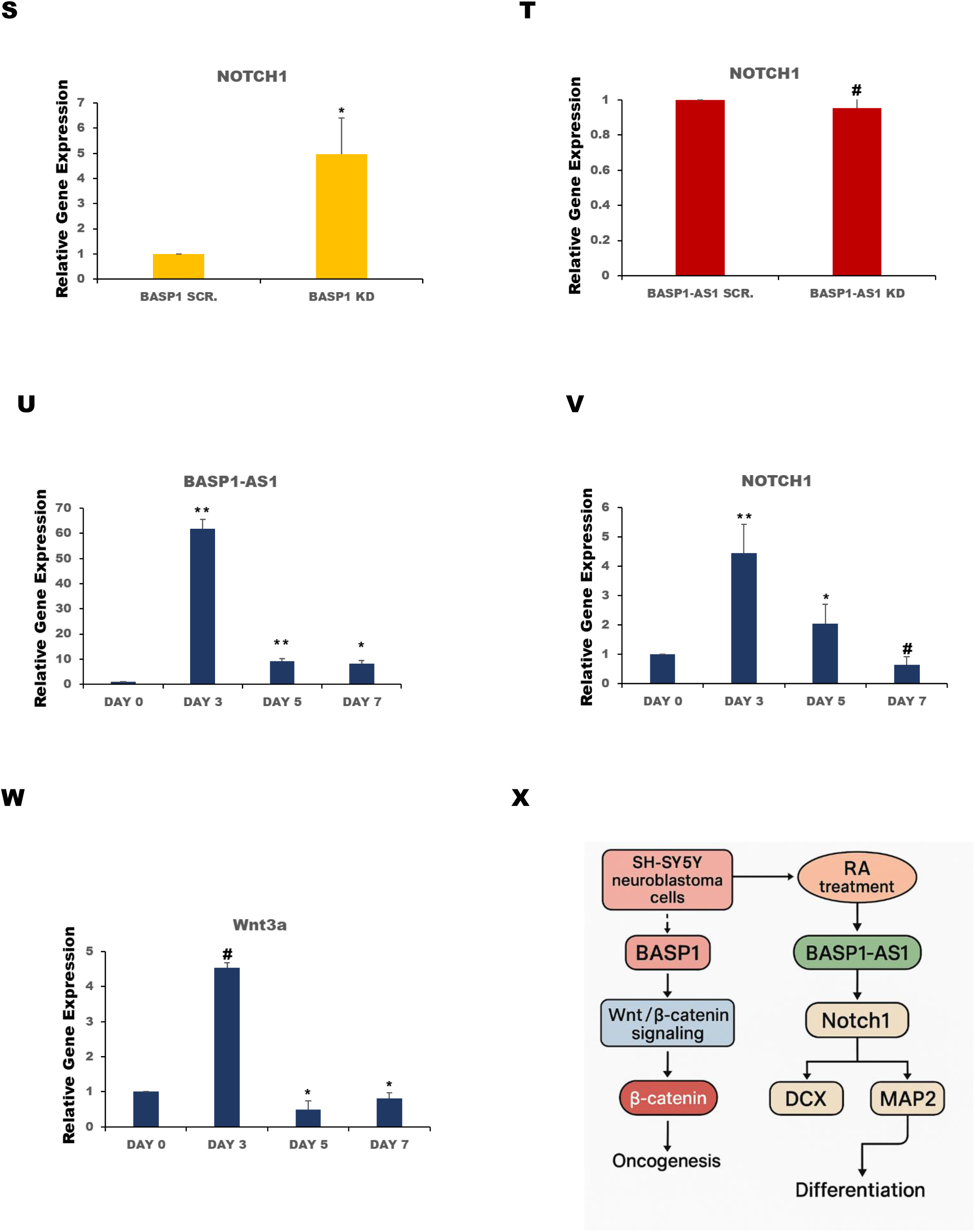
| Reciprocal regulation between BASP1 and BASP1-AS1 modulates Notch1 signaling and promotes neuronal marker expression. (A–E) Western blot analysis showing early progressive decline in β-catenin, DCX, and Sox2 protein levels during RA-induced differentiation, maintained at low levels throughout the time course. (F) Morphological changes observed in SH-SY5Y cells transfected with BASP1 siRNA and exposed to differentiation medium, showing elongated somas and extensive neurite outgrowth after 72 hours. (G–H) BASP1 knockdown leads to significant upregulation of BASP1-AS1 lncRNA expression. (I) Schematic of the regulatory feedback loop between BASP1 and BASP1-AS1 under RA treatment. BASP1-AS1 promotes BASP1 expression under basal conditions; RA-induced BASP1 knockdown upregulates BASP1-AS1, maintaining a regulatory balance. (J–K) BASP1-AS1 expression positively correlates with increased DCX levels, which peak at day 3 and remain elevated throughout differentiation. (L–M) Knockdown of BASP1-AS1 suppresses DCX expression, supporting its role in DCX transcriptional regulation. (N–O) BASP1-AS1 expression positively correlates with MAP2 levels, whereas its knockdown reduces MAP2 expression, indicating BASP1-AS1 is sufficient but not essential for MAP2 induction. (P–Q) Elevated BASP1-AS1 has no significant impact on Sox2 or Ki67 expression. (R–S) BASP1 knockdown results in reduced Wnt3a and increased Notch1 expression. (T) BASP1-AS1 knockdown does not affect Notch1 expression, indicating Notch1 upregulation is specifically associated with BASP1-AS1 elevation. (U) Time-course analysis of BASP1-AS1 expression reveals a transient 60-fold increase at day 3 post-RA treatment, followed by a decline and stabilization above baseline. (V) Notch1 shows similar temporal expression dynamics, peaking at day 3 and declining thereafter. (W) Wnt3a expression shows a non-significant 4-fold increase at day 3, followed by a progressive decline to 50% and 20% of baseline by days 5 and 7, respectively. (X) Schematic model summarizing the regulatory cascade linking BASP1 knockdown and BASP1-AS1 upregulation to suppression of Wnt/β-catenin signaling and activation of Notch1-driven neuronal differentiation. Solid arrows indicate activation or expression relationships; specifically, the arrow from SH-SY5Y cells to BASP1 denotes endogenous expression of BASP1 in this cell line. All data represent mean ± SD from at least three independent experiments. Statistical significance was determined by Student’s t-test. *P < 0.05, ** P < 0.005, ** P < 0.0005 ^#^ p ≥ 0.05

BASP1 knockdown decreased proliferating and differentiating markers in undifferentiated SH-SY5Y cells. To assess the role of BASP1 in differentiating cells, undifferentiated SH-SY5Y cells transfected with BASP1 specific siRNA under standard conditions exhibited distinct morphological changes within 72 hours of culture in differentiation-specific media. Morphologically, BASP1 knockdown cells displayed elongated somas, extensive neurite outgrowth, and a more complex neuronal architecture (Figure 3F).

BASP1 knockdown led to a significant upregulation of BASP1-AS1 lncRNA expression (Figure 3G-H). A regulatory pathway diagram to visually represent the regulatory relationships between BASP1-AS1 and BASP1 has been shown in Figure 3I. The diagram illustrates the regulatory interplay between the long non-coding RNA BASP1-AS1 and its target gene BASP1. In the absence of retinoic acid, downregulation of BASP1-AS1 leads to reduced expression of BASP1, indicating a positive regulatory role. Upon retinoic acid treatment, the depletion of BASP1 in turn activates BASP1-AS1, forming a feedback loop. This loop suggests that retinoic acid indirectly maintains homeostasis between the two transcripts through reciprocal regulation. Upregulation of BASP1-AS1 lncRNA expression correlated with increased DCX transcript levels (Figure 3J). DCX expression peaked markedly on day 3 and, although levels declined thereafter, they remained approximately five-fold higher than in undifferentiated cells throughout the observation window (Figure 3K). Conversely, knockdown of BASP1-AS1 led to a suppression of DCX expression (Figure 3L-M), supporting a regulatory role for BASP1-AS1 in driving DCX expression. Upregulation of BASP1-AS1 enhances MAP2 expression, whereas BASP1-AS1 knockdown lowered MAP2 levels (Figure 3N-O). This indicates that BASP1-AS1 is sufficient, but not essential, for MAP2 upregulation. Thus, while increased BASP1-AS1 can promote MAP2 expression, MAP2 regulation can also occur through alternative pathways in the absence of BASP1-AS1, suggesting the presence of compensatory mechanisms that maintain MAP2 expression during neuronal differentiation. Thus, BASP1-AS1 upregulation is sufficient to induce neuronal maturation markers, supporting its role as a positive regulator of neuronal differentiation. Notably, elevated BASP1-AS1 levels following BASP1 knockdown had no measurable impact on Sox2 or Ki67 expression (Figure 3P-Q).

Knockdown of BASP1, which led to an upregulation of BASP1-AS1, was associated with a reduction in Wnt3a expression and an increase in Notch1 expression (Figure 3R-S). In contrast, BASP1-AS1 knockdown had no significant effect on Notch1 levels (Figure 3T), indicating that Notch1 activation is specifically associated with elevated BASP1-AS1. Given the known role of Notch signaling in promoting neuroblastoma cells differentiation [29, 30], these findings suggest that transient upregulation of BASP1-AS1 relieves repression by BASP1, leading to transcriptional activation of Notch1 and differentiation associated pathways.

BASP1-AS1 expression remained low in undifferentiated SH-SY5Y cells. Upon RA-induced differentiation, a sharp and transient increase in BASP1-AS1 expression was observed, peaking around day 3 with an approximately 60-fold elevation relative to undifferentiated cells (Figure 3U). This peak was followed by a gradual decline, with expression levels stabilizing at approximately 8-fold higher than baseline by days 5 and 7. These findings suggest BASP1-AS1 is tightly regulated and temporally coordinated with early differentiation events. Like BASP1-AS1, Notch1 exhibited a peak at day 3 before declining on days 5 and 7 (Figure 3V). Expression levels of Wnt3a declined by 50% and 20%, by day 5 and day 7 respectively (Figure 3W).

Our model illustrates a shift from oncogenic Wnt/β-catenin signaling toward a differentiation-promoting Notch1 pathway, coordinated through BASP1-AS1 expression (Figure 3X). The upregulation of BASP1-AS1 activates Notch1 expression, a key regulator of neuronal differentiation. Activated Notch1 signaling then enhances the expression of neuronal markers DCX (Doublecortin) and MAP2 (Microtubule-Associated Protein 2), both critical for the acquisition of a differentiated neuronal phenotype. This pathway highlights the dynamic regulatory interplay during RA-induced differentiation, where BASP1-AS1 acts as a molecular switch linking the suppression of Wnt3a–driven oncogenesis to the activation of Notch1-mediated differentiation. The temporal expression of BASP1-AS1 thus serves as a regulatory hub in ensuring proper lineage commitment in neuroblastoma cells.

### 4. BASP1-AS1 regulates a positive feedback loop involving BASP1 and non-canonical Wnt2 signaling during Neuronal Differentiation

BASP1 expression showed a modest transient induction-peaking at ∼60% above baseline on day 3 – before decreasing below undifferentiated levels by day 7 (Figure 4A). Specifically, Wnt2 showed an approximately 20-fold increase over baseline at day 3, which then dropped sharply by day 5 and partially rebounded to about twofold above basal levels by day 7 (Figure 4B).

**Figure 4.**
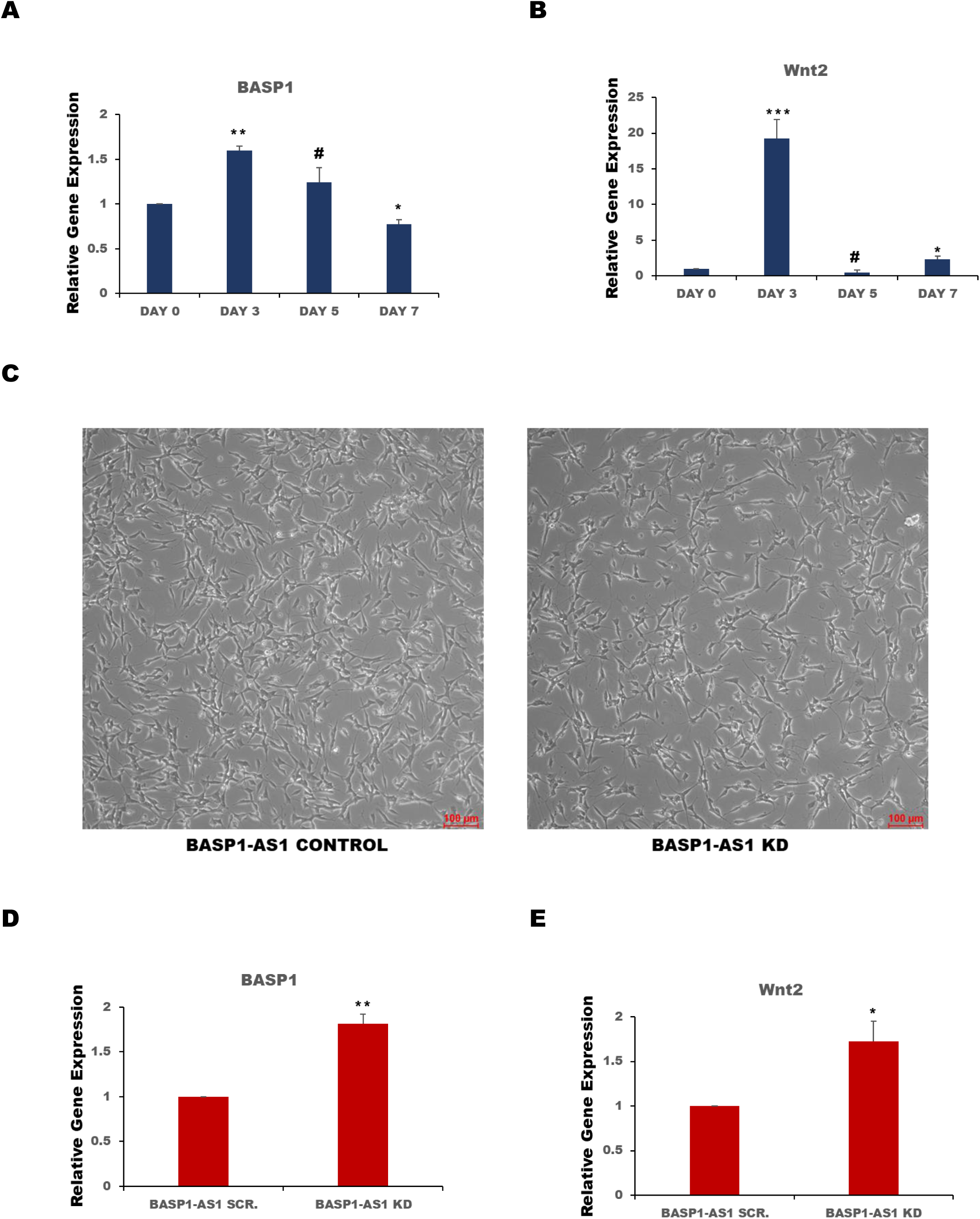
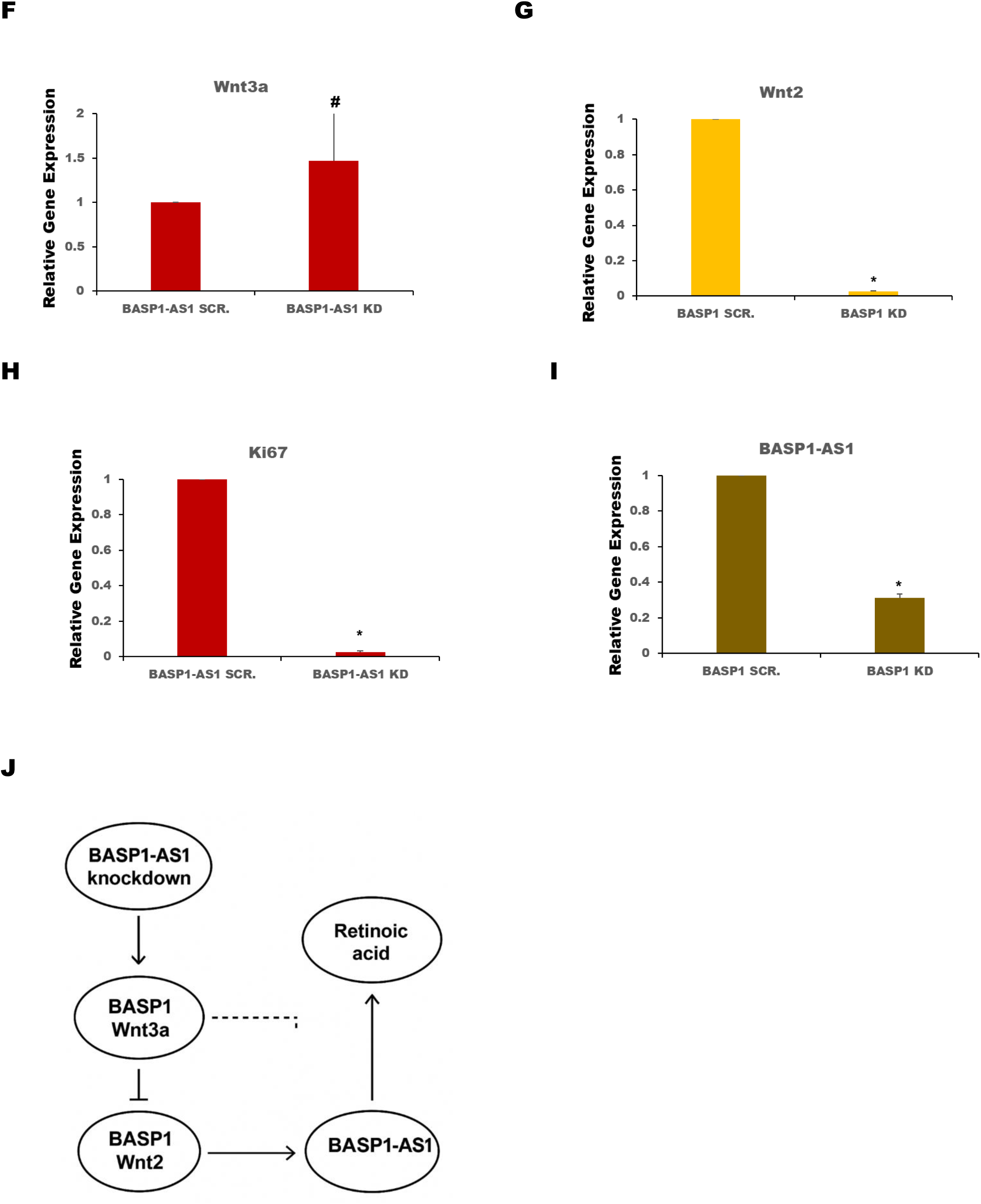
| BASP1-AS1 regulates a positive feedback loop involving BASP1 and non-canonical Wnt2 signaling during differentiation. (A) BASP1 mRNA expression exhibits a transient increase, peaking at ∼60% above undifferentiated levels on day 3, followed by a decrease below baseline by day 7. (B) Wnt2 expression shows a ∼20-fold induction at day 3 post-RA treatment, declining sharply by day 5 and stabilizing at ∼2-fold above baseline by day 7. (C) Morphological analysis of undifferentiated SH-SY5Y cells transfected with BASP1-AS1 siRNA and incubated in differentiation medium for 72 hours reveals accelerated neurite outgrowth, enhanced branching, and reduced proliferative morphology. (D–E) Silencing of BASP1-AS1 results in robust upregulation of BASP1 (D) and Wnt2 (E)mRNA expression. (F) Wnt3a expression remains unchanged upon BASP1-AS1 knockdown, suggesting selective engagement of Wnt signaling components. (G) BASP1 knockdown leads to decreased Wnt2 expression, reinforcing the dependence of Wnt2 on BASP1 activity. (H) Induction of the BASP1/Wnt2 axis following BASP1-AS1 knockdown is accompanied by decreased Ki67 expression, indicating reduced proliferation and promotion of differentiation. (I) Silencing of BASP1 on day 1 of RA treatment leads to decreased BASP1-AS1 expression, indicating a positive regulatory effect of the BASP1/Wnt2 axis on BASP1-AS1. (J) Schematic model depicting the context-dependent regulatory interactions among BASP1-AS1, BASP1, Wnt2/Wnt3a, and retinoic acid. In the absence of RA, BASP1-AS1 suppresses BASP1 and Wnt3a, limiting Wnt2 expression. Under RA treatment, BASP1-AS1 knockdown activates BASP1 and Wnt2, which in turn upregulate BASP1-AS1, forming a positive feedback loop that promotes differentiation. Solid arrows represent activation, dashed arrows represent inhibition, and ovals denote molecular entities. All data represent mean ± SD from at least three independent experiments. Statistical significance was determined by Student’s t-test. *P < 0.05, ** P < 0.005, ** P < 0.0005 ^#^ p ≥ 0.05

BASP1-AS1 decreased the BASP1 expression in undifferentiated cells to reduce cellular proliferation. So, SH-SY5Y cells were transfected with BASP1-AS1 specific siRNA under standard conditions and were incubated for 72 hours in differentiation-specific media. Knockdown (KD) cells displayed accelerated neurite outgrowth and branching, forming complex intercellular connections. Compared to control cultures, BASP1-AS1-depleted cells showed reduced proliferative morphology, consistent with a phenotypic shift toward neuronal lineage commitment (Figure 4C). Silencing of BASP1-AS1 resulted in a robust upregulation of BASP1 expression (Figure 4D), accompanied by increased Wnt2 transcript levels (Figure 4E). In contrast, Wnt3a expression remained unchanged (Figure 4F), indicating selective engagement of Wnt signaling components in differentiation specific media. Knockdown of BASP1 suppressed Wnt2 expression (Figure 4G). BASP1/Wnt2 mRNA induction following knockdown of BASP1-AS1 was accompanied by a notable reduction in Ki67 gene expression, suggesting a decline in proliferative activity and reinforcing the onset of neuronal differentiation (Figure 4H). Silencing of BASP1 using siRNA on day 1 of differentiation led to a decreased expression of BASP1-AS1 (Figure 4I). This suggests that following the initiation of differentiation, activation of the BASP1/Wnt2 axis positively regulates BASP1-AS1 expression. A regulatory pathway diagram of BASP1-AS1, BASP1/Wnt signaling, and retinoic acid has been shown in Figure 4J. In the absence of retinoic acid, BASP1-AS1 knockdown leads to downregulation of BASP1 and Wnt3a, resulting in suppressed Wnt2 expression. In contrast, under retinoic acid treatment, BASP1-AS1 knockdown increases BASP1 and Wnt2 expression. The elevated BASP1/Wnt2 axis then enhances BASP1-AS1 expression, forming a positive feedback loop.

### 5. Canonical Wnt Activation by LiCl Represses Differentiation through repression of BASP1-AS1

SH-SY5Y neuroblastoma cells treated with the Wnt pathway activator lithium chloride (LiCl) in RA differentiation-specific media exhibited a marked downregulation of BASP1-AS1 transcripts (Figure 5A). Wnt3a expression increased following LiCl treatment, consistent with the activation of canonical Wnt/β-catenin signaling (Figure 5B). This was further supported by enhanced levels of active β-catenin protein. Notably, LiCl treatment also led to a robust upregulation of Sox2 protein expression (Figure 5C-E), a transcription factor central to the maintenance of neuronal plasticity and stemness. These findings collectively indicate that LiCl-induced activation of Wnt/β-catenin signaling suppresses BASP1-AS1, thereby contributing to the emergence of a stem-like, therapy-resistant subpopulation within neuroblastoma cultures.

**Figure 5.**
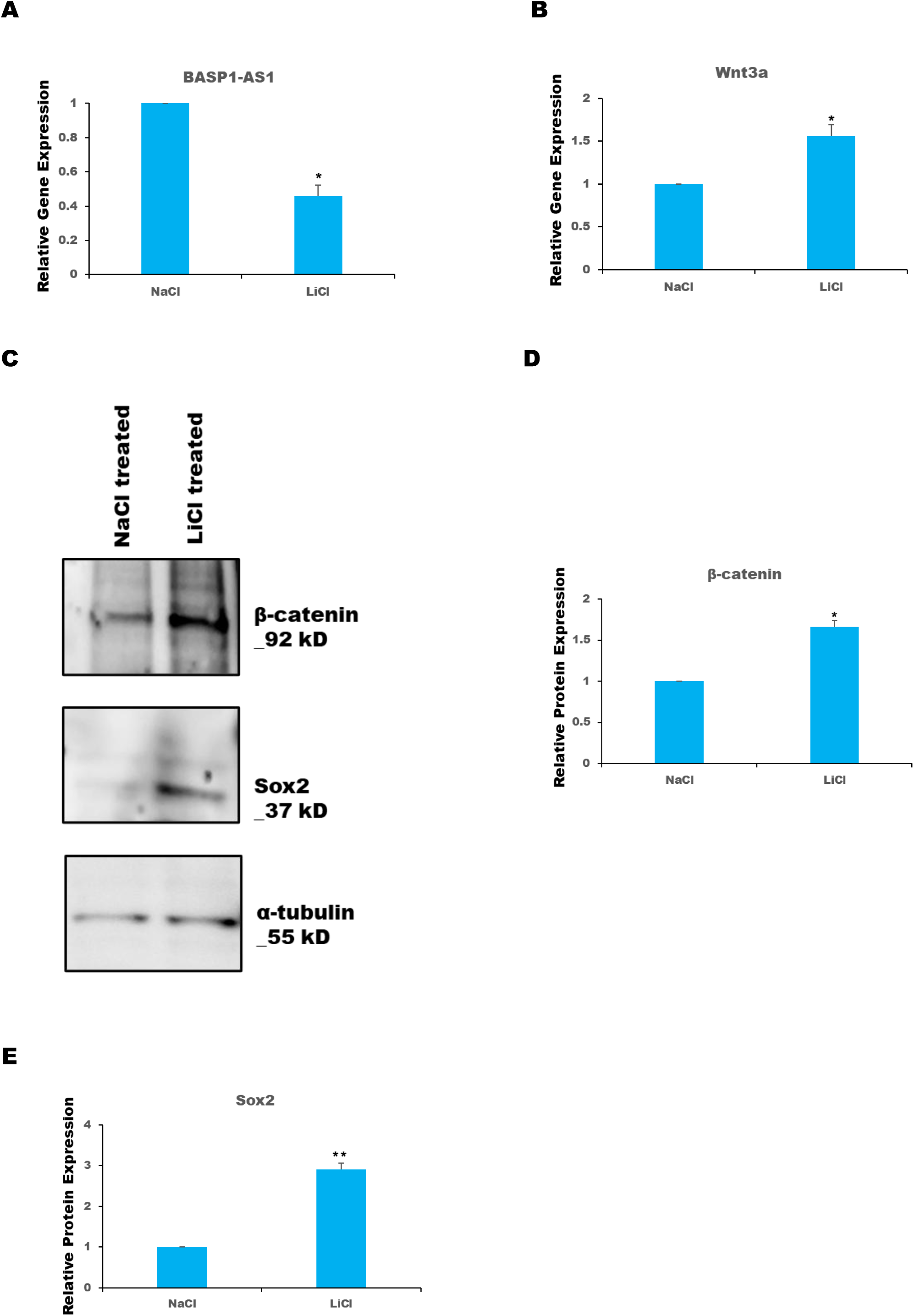
| Canonical Wnt activation by LiCl suppresses neuronal differentiation through repression of BASP1-AS1. (A) Treatment of SH-SY5Y cells with LiCl in RA differentiation medium significantly downregulates BASP1-AS1 transcript levels. (B) LiCl treatment induces Wnt3a mRNA expression, consistent with activation of canonical Wnt/β-catenin signaling. (C–E) Immunoblot analysis reveals increased levels of active β-catenin and robust upregulation of Sox2 protein in LiCl-treated cultures. All data represent mean ± SD from at least three independent experiments. Statistical significance was determined by Student’s t-test. *P < 0.05.

### 6. Canonical Wnt Inhibition by DKK1 Promotes Differentiation via Activation of BASP1-AS1

SH-SY5Y neuroblastoma cells treated with the Wnt pathway inhibitor dickopff1 (DKK1) in RA differentiation-specific media, increased BASP1-AS1 transcripts (Figure 6A). DKK1 also increased neuronal DCX and MAP2 gene expression, suggesting enhanced differentiation (Figure 6B-C). Additionally, DKK1 treatment led to a downregulation of Wnt3a expression levels (Figure 6D). Notably, increased levels of BASP1-AS1, in the context of reduced Wnt3a expression, may be more favourable compared to a cellular state characterized by low BASP1-AS1 and high Wnt3a expression.

**Figure 6.**
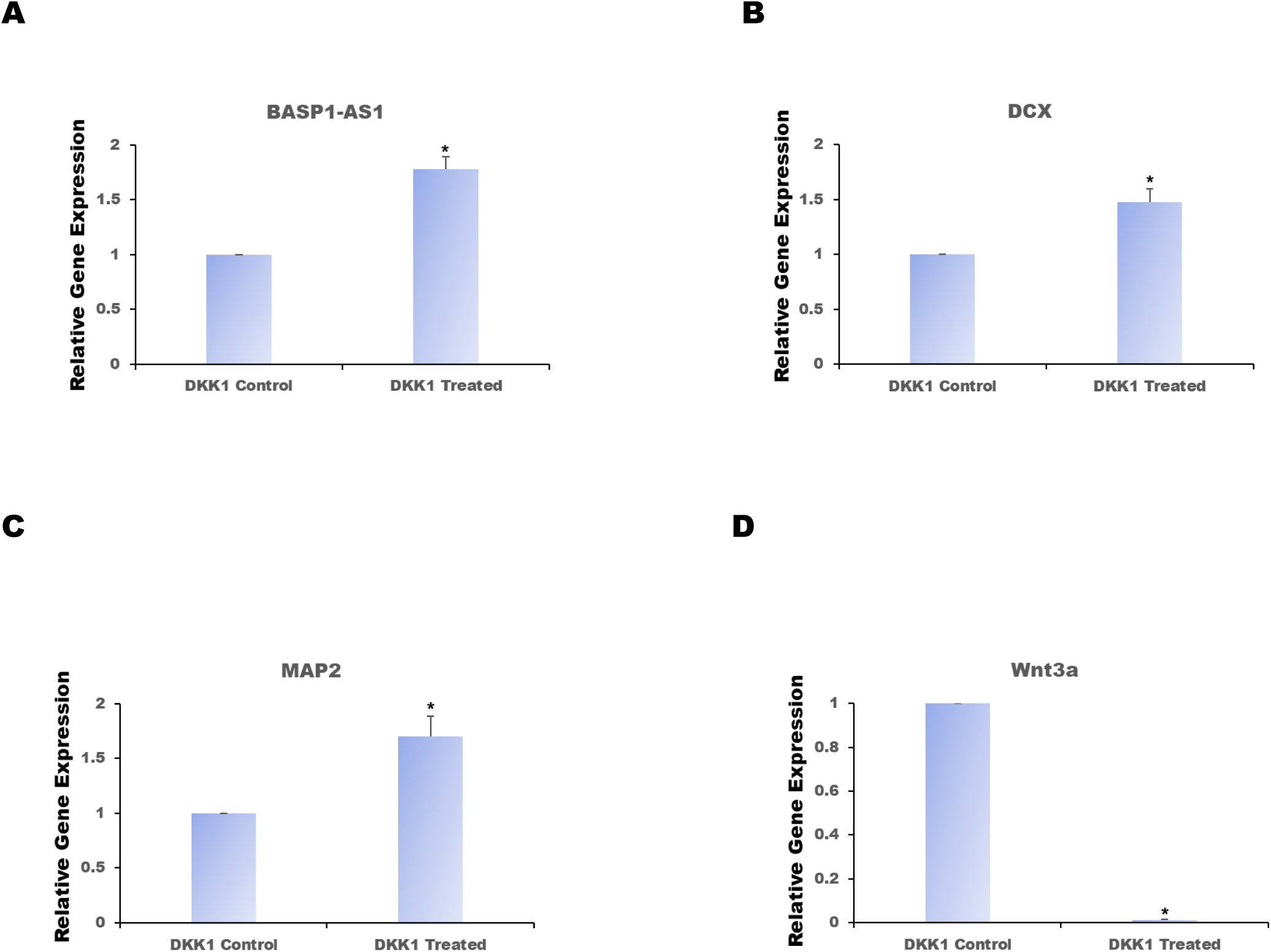
| Canonical Wnt inhibition by DKK1 promotes neuronal differentiation via activation of BASP1-AS1. (A) Treatment of SH-SY5Y neuroblastoma cells with DKK1 in RA differentiation medium significantly upregulates BASP1-AS1 transcript levels. (B–C) DKK1 treatment increases expression of neuronal differentiation markers DCX (B) and MAP2 (C), indicating enhanced neurogenic progression. (D) Wnt3a mRNA expression is significantly reduced following DKK1 treatment. All data represent mean ± SD from at least three independent experiments. Statistical significance was determined by Student’s t-test. *P < 0.05.

Upon Wnt activation in RA-treated cells, Wnt3a expression increases. Activation of the Wnt signaling pathway in RA-treated conditions leads to a reduction in BASP1-AS1 expression, which may contribute to therapeutic resistance. This creates a complex molecular landscape characterized by low BASP1-AS1, and elevated Wnt3a levels – an expression profile that may be associated with a less favourable therapeutic response.

### 7. BDNF Defines Transcriptional Thresholds for Terminal Neuronal Maturation in SH-SY5Y Cells

To define the threshold expression levels of key differentiation regulators during terminal neuronal maturation, SH-SY5Y neuroblastoma cells were treated with retinoic acid (RA) for 5 days to initiate differentiation and subsequently exposed to brain-derived neurotrophic factor (BDNF) in serum-free medium till 13^th^ day (Figure 7A). Gene expression analysis was performed to compare mature, BDNF-treated neurons with undifferentiated neuroblastoma cells.

**Figure 7.**
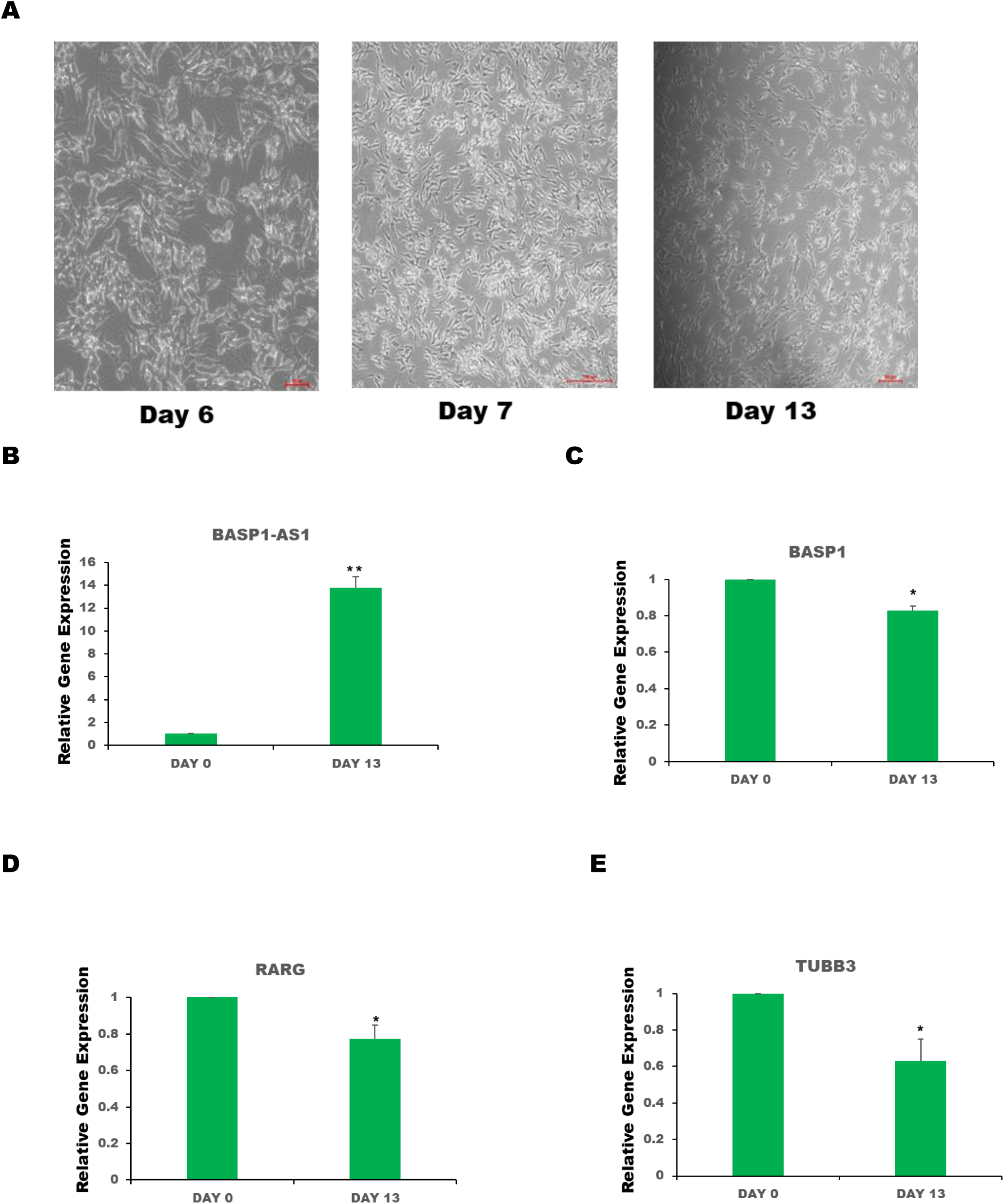
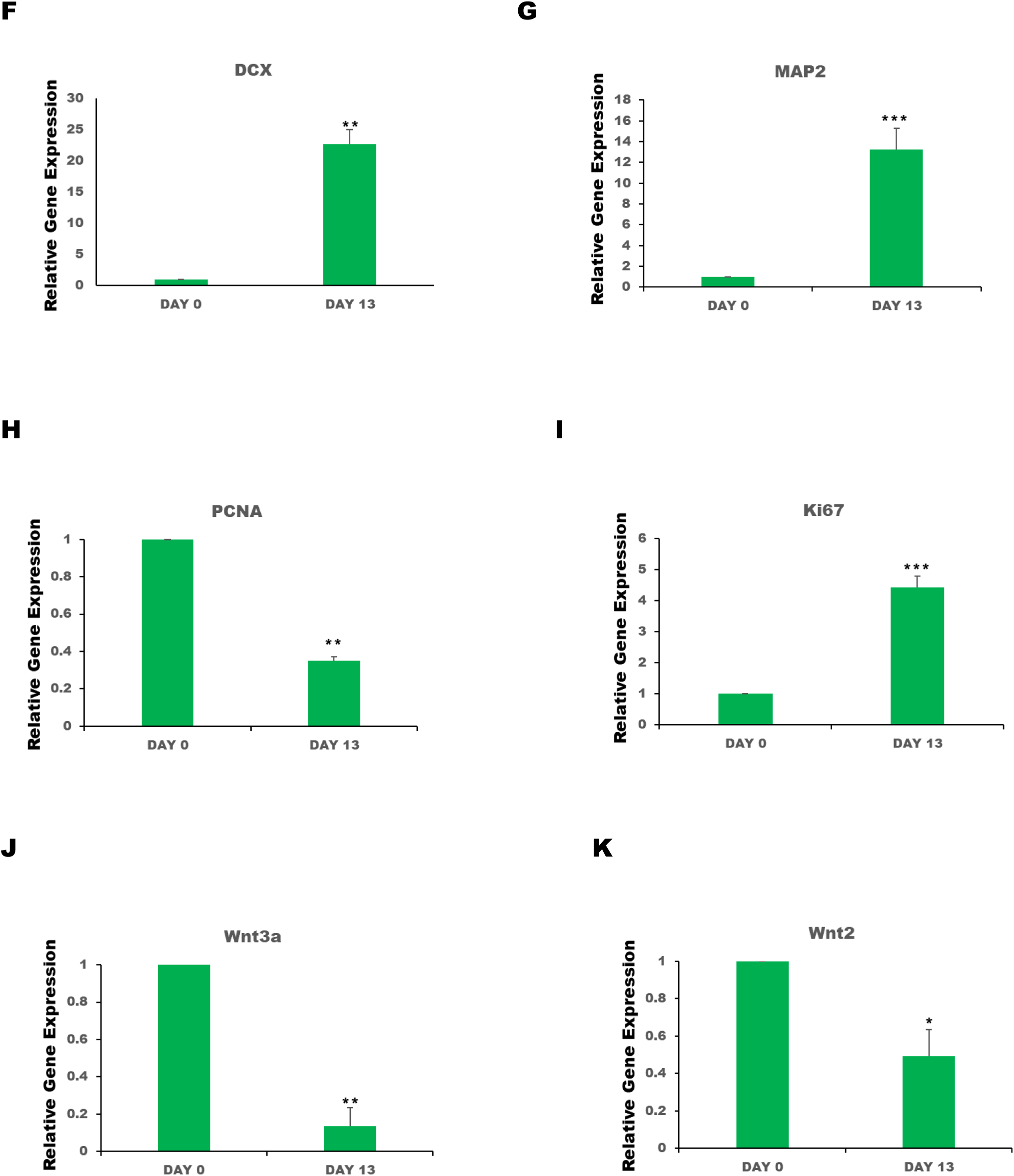
| BDNF defines transcriptional thresholds for terminal neuronal maturation in SH-SY5Y cells. (A) SH-SY5Y cells were treated with RA for 5 days, followed by BDNF exposure to induce terminal maturation. (B) BASP1-AS1 expression is significantly upregulated in BDNF-treated neurons, suggesting a role in stabilizing the mature neuronal phenotype. (C) BASP1 mRNA levels are markedly reduced following BDNF treatment, indicating its repression during terminal differentiation. (D–E) Expression of early differentiation markers RARG and TUBB3 is significantly downregulated in BDNF-matured neurons relative to undifferentiated cells. (F–G) Terminal neuronal markers DCX and MAP2 are robustly upregulated in response to BDNF, consistent with enhanced neuronal migration and maturation. (H) PCNA expression is significantly decreased in mature neurons, reflecting cell cycle exit. (I) Ki67 remains detectable at low levels, suggesting either residual proliferation or cell cycle-independent functions. (J–K) BDNF treatment leads to strong downregulation of Wnt3a (∼90%) and Wnt2 (∼50%), in terminally differentiated cells. All data represent mean ± SD from at least three independent experiments. Statistical significance was determined by Student’s t-test. *P < 0.05, ** P < 0.005, ** P < 0.0005

BASP1-AS1 expression was significantly upregulated in BDNF-treated cells (Figure 7B), suggesting its involvement in maintaining mature neuronal identity. In contrast, BASP1 expression was markedly decreased (Figure 7C), despite its established role in early differentiation, indicating that its function may be restricted to the early neurogenic commitment phase and actively repressed during maturation.

Consistent with the transition from early differentiation to terminal maturation, the expression of RARG and TUBB3-both associated with neuronal commitment-was significantly downregulated in BDNF-treated neurons relative to undifferentiated controls (Figure 7D-E). This downregulation marks the resolution of progenitor-like programs and the cessation of early differentiation signals.

Markers of neuronal migration and maturation were substantially elevated. DCX expression increased approximately 22-fold, while MAP2 levels rose 13-fold in response to BDNF treatment (Figure 7F-G), confirming progression toward a migratory, structurally mature neuronal phenotype.

Cell proliferation markers exhibited a distinct and divergent expression pattern. PCNA expression was significantly reduced in mature neurons, consistent with terminal cell cycle exit (Figure 7H). In contrast, Ki67 expression remained detectable (Figure 7I), suggesting that a minor subpopulation may retain low-level proliferative capacity or that Ki67 may exert cell cycle–independent functions in post-mitotic neurons.

BDNF-induced maturation was also characterized by suppression of Wnt signaling. Expression of Wnt3a was reduced by ∼90%, and Wnt2 levels declined by ∼50% relative to undifferentiated progenitors (Figure 7J-K). This suggests that attenuation of Wnt signaling is a hallmark of terminal neuronal differentiation.

Collectively, these results highlight a transcriptional threshold pattern in which high BASP1-AS1 and DCX/MAP2 expression, coupled with downregulation of BASP1, Wnt components, and proliferation markers, defines the terminally differentiated neuronal state. The discordance between PCNA and Ki67 expression further implies heterogeneity within the mature population and reflects transitional or non-canonical roles of proliferation-associated genes during neuronal maturation.

## Discussion

Neuroblastoma remains one of the most challenging paediatric malignancies due to its high intra-tumoral heterogeneity and therapy-resistant subpopulations. Our study identifies BASP1 and its antisense lncRNA, BASP1-AS1, as critical regulators of proliferation and differentiation in SH-SY5Y neuroblastoma cells. We uncover a dynamic interplay between canonical and non-canonical Wnt signaling pathways, orchestrated by a BASP1/BASP1-AS1 regulatory axis, that determines lineage commitment and neuronal maturation. This axis functions as a molecular switch, facilitating the transition from a proliferative, Wnt3a-driven oncogenic state to a differentiated, Notch1-mediated neuronal phenotype.

### BASP1 Acts as a Context-Dependent Regulator of Proliferation and Differentiation

Our findings highlight the dual roles of BASP1 in SH-SY5Y cells: it supports proliferative capacity via Wnt3a signaling in undifferentiated states, while its downregulation is required for neuronal differentiation. Knockdown of BASP1 led to reduced expression of proliferation markers (Ki67, PCNA) and neuronal markers (DCX, MAP2), indicating a critical role in maintaining a poised, progenitor-like state. Importantly, BASP1 silencing also reduced Wnt3a expression, implicating it as a downstream effector of Wnt/β-catenin signaling. These data position BASP1 as a gatekeeper gene, maintaining oncogenic signaling under homeostatic conditions while restraining premature differentiation.

### BASP1-AS1 Functions as a Differentiation-Activating lncRNA

The long non-coding RNA BASP1-AS1 emerged as a pivotal positive regulator of neuronal fate. RA-induced differentiation resulted in a rapid and transient upregulation of BASP1-AS1, temporally coinciding with the downregulation of Sox2 and β-catenin. BASP1-AS1 depletion impaired DCX upregulation and attenuated MAP2 induction. These findings suggest BASP1-AS1 acts as an early-response lncRNA to RA and plays a key role in the commitment to neuronal lineages.

### A Molecular Switch from Wnt3a to Notch1 Signaling

Our results support a model where RA induces a transcriptional switch from canonical Wnt signaling (via Wnt3a and β-catenin) to Notch1 activation, mediated by BASP1-AS1. Notch1 upregulation upon BASP1 knockdown and BASP1-AS1 induction – without reciprocal activation upon BASP1-AS1 silencing – suggests that BASP1-AS1 serves as a necessary intermediate in disinhibiting Notch1 signaling. Given Notch1’s role in promoting differentiation [29,30], this axis may represent a tightly regulated checkpoint, ensuring that proliferation-associated signaling (β-catenin, Sox2) is silenced before differentiation ensues.

Interestingly, treatment with the canonical Wnt agonist LiCl repressed BASP1-AS1 expression, and activated β-catenin, thereby reinforcing a proliferative, stem-like phenotype characterized by elevated Sox2 expression. This inverse relationship between canonical Wnt activation and BASP1-AS1 expression emphasizes the antagonistic regulation of lineage commitment between them.

### BASP1-AS1/Wnt2 Axis Engages Non-Canonical Differentiation Pathways

Our data suggest that in the presence of RA, the BASP1-AS1/BASP1/Wnt2 axis may activate a β-catenin–independent, non-canonical Wnt pathway that promotes differentiation. This is supported by the selective induction of Wnt2 – but not Wnt3a – upon BASP1-AS1 knockdown, along with decreased Ki67 expression and enhanced neuronal morphology. Importantly, BASP1 silencing inhibited Wnt2 induction, indicating that Wnt2 lies downstream of BASP1. These findings highlight a positive feedback loop, wherein Wnt2 signaling maintains BASP1-AS1 expression, further reinforcing differentiation. Taken together, our data suggest that targeting the Wnt pathway – specifically Wnt2 activation may enhance pro-differentiation programs via BASP1-AS1 upregulation, offering a new avenue for overcoming differentiation blockades in high-risk or RA-refractory neuroblastoma. This non-canonical Wnt axis may represent a compensatory or alternative differentiation pathway that bypasses the oncogenic β-catenin circuit, with implications for targeting neuroblastoma subtypes that are resistant to RA-induced maturation.

### DKK1-Mediated Wnt Suppression Reveals BASP1-AS1 as a Regulatory Hub in Neuroblastoma Differentiation

Our findings reveal that pharmacological inhibition of β-catenin signaling via DKK1 induces an enhanced neuronal differentiation program in RA treated SH-SY5Y neuroblastoma cells, characterized by upregulation of DCX and MAP2. Intriguingly, DKK1-mediated downregulation of Wnt3a correlates with increased BASP1-AS1 expression, indicating that repression of Wnt3a may be a critical node influencing this axis. Notably, the observed elevation of BASP1-AS1 in the presence of low Wnt3a may represent a more therapeutically favourable transcriptional state, particularly when contrasted with RA-treated cells, where Wnt activation leads to high Wnt3a, and suppressed BASP1-AS1 levels – an expression signature potentially linked to resistance.

### BDNF as a Terminal Differentiation Cue Defines a New Transcriptional Threshold

Upon BDNF treatment, BASP1-AS1 expression was dramatically upregulated while BASP1 was repressed, indicating a biphasic model in which BASP1 is essential for early differentiation but must be silenced for terminal maturation. This stage-specific repression was accompanied by elevated DCX and MAP2, along with near-complete suppression of Wnt3a and significant downregulation of Wnt2, confirming that attenuation of both Wnt pathways is required for stable neuronal identity.

Interestingly, while PCNA levels were reduced, Ki67 remained detectable, suggesting possible cell-cycle–independent roles for Ki67 or the presence of a transiently cycling subpopulation. This observation aligns with recent reports describing Ki67 expression in terminally differentiated cells, where it may regulate chromatin organization rather than proliferation [31,32].

### Therapeutic Implications and Future Directions

This study offers significant implications for differentiation therapy in neuroblastoma. Targeting BASP1 or modulating the BASP1-AS1/Notch1 axis could enable therapeutic rewiring of proliferative neuroblastoma cells toward differentiation. Moreover, canonical Wnt activation may serve as a biomarker of resistance or dedifferentiation, especially in the context of RA-based therapies. BASP1-AS1, by mediating the shift from oncogenic to differentiation pathways, emerges as a potential lncRNA therapeutic, warranting further investigation for delivery or activation strategies.

In conclusion, our findings delineate a novel lncRNA-mediated switch that coordinates the balance between proliferation and differentiation in neuroblastoma. The BASP1/BASP1-AS1 axis represents a dynamic regulatory module, essential for neurogenic lineage progression, and provides a foundation for developing lncRNA-targeted differentiation therapies in paediatric cancer.

## Author contributions

S.K: Conceptualized and designed the study, Performed experiments, Data analysis and interpretation. Wrote the manuscript. M.K: Performed experiments, Data analysis

## References

1. Maris JM, Hogarty MD, Bagatell R, Cohn SL. Neuroblastoma. Lancet. 2007

2. Brodeur GM. Neuroblastoma: biological insights into a clinical enigma. Nat Rev Cancer. 2003

3. Cohn SL, Pearson AD, London WB, Monclair T, Ambros PF, Brodeur GM, Faldum A, Hero B, Iehara T, Machin D, Mosseri V, Simon T, Garaventa A, Castel V, Matthay KK; INRG Task Force. The International Neuroblastoma Risk Group (INRG) classification system: an INRG Task Force report. J Clin Oncol. 2009

4. Park JR, Bagatell R, London WB, Maris JM, Cohn SL, Mattay KK, Hogarty M; COG Neuroblastoma Committee. Children’s Oncology Group’s 2013 blueprint for research: neuroblastoma. Pediatr Blood Cancer. 2013

5. Chabner BA, Roberts TG Jr. Timeline: Chemotherapy and the war on cancer. Nat Rev Cancer. 2005

6. Matthay KK, Reynolds CP, Seeger RC, Shimada H, Adkins ES, Haas-Kogan D, Gerbing RB, London WB, Villablanca JG. Long-term results for children with high-risk neuroblastoma treated on a randomized trial of myeloablative therapy followed by 13-cis-retinoic acid: a children’s oncology group study. J Clin Oncol. 2009

7. Sidell N. Retinoic acid-induced growth inhibition and morphologic differentiation of human neuroblastoma cells in vitro. J Natl Cancer Inst. 1982

8. Reynolds CP, Matthay KK, Villablanca JG, Maurer BJ. Retinoid therapy of high-risk neuroblastoma. Cancer Lett. 2003

9. Altucci L, Gronemeyer H. The promise of retinoids to fight against cancer. Nat Rev Cancer. 2001

10. Nusse R, Clevers H. Wnt/β-Catenin Signaling, Disease, and Emerging Therapeutic Modalities. Cell. 2017

11. Yang Y. Wnt signaling in development and disease. Cell Biosci. 2012

12. Duffy DJ, Krstic A, Schwarzl T, Halasz M, Iljin K, Fey D, Haley B, Whilde J, Haapa-Paananen S, Fey V, Fischer M, Westermann F, Henrich KO, Bannert S, Higgins DG, Kolch W. Wnt signalling is a bi-directional vulnerability of cancer cells. Oncotarget. 2016

13. Szemes M, Greenhough A, Malik K. Wnt Signaling Is a Major Determinant of Neuroblastoma Cell Lineages. Front Mol Neurosci. 2019

14. Flahaut M, Meier R, Coulon A, Nardou KA, Niggli FK, Martinet D, Beckmann JS, Joseph JM, Mühlethaler-Mottet A, Gross N. The Wnt receptor FZD1 mediates chemoresistance in neuroblastoma through activation of the Wnt/beta-catenin pathway. Oncogene. 2009

15. Zhi F, Gong G, Xu Y, Zhu Y, Hu D, Yang Y, Hu Y. Activated β-catenin forces N2A cell-derived neurons back to tumor-like neuroblasts and positively correlates with a risk for human neuroblastoma. Int J Biol Sci. 2012

16. Krishna S, Prajapati B, Seth P, Sinha S. Dickopff 1 inhibits cancer stem cell properties and promotes neuronal differentiation of human neuroblastoma cell line SH-SY5Y. IBRO Neurosci Rep. 2024

17. Zins K, Schäfer R, Paulus P, Dobler S, Fakhari N, Sioud M, Aharinejad S, Abraham D. Frizzled2 signaling regulates growth of high-risk neuroblastomas by interfering with β-catenin-dependent and β-catenin-independent signaling pathways. Oncotarget. 2016

18. Duffy DJ, Krstic A, Schwarzl T, Halasz M, Iljin K, Fey D, Haley B, Whilde J, Haapa-Paananen S, Fey V, Fischer M, Westermann F, Henrich KO, Bannert S, Higgins DG, Kolch W. Wnt signalling is a bi-directional vulnerability of cancer cells. Oncotarget. 2016

19. Becker J, Wilting J. WNT Signaling in Neuroblastoma. Cancers (Basel). 2019

20. Li C, Song G, Zhang S, Wang E, Cui Z. Wnt3a increases the metastatic potential of non-small cell lung cancer cells in vitro in part via its upregulation of Notch3. Oncol Rep. 2015

21. Nwabo Kamdje AH, Takam Kamga P, Tagne Simo R, Vecchio L, Seke Etet PF, Muller JM, Bassi G, Lukong E, Kumar Goel R, Mbo Amvene J, Krampera M. Developmental pathways associated with cancer metastasis: Notch, Wnt, and Hedgehog. Cancer Biol Med. 2017

22. Brodeur GM. Neuroblastoma: biological insights into a clinical enigma. Nat Rev Cancer. 2003

23. Agholme L, Lindström T, Kågedal K, Marcusson J, Hallbeck M. An in vitro model for neuroscience: differentiation of SH-SY5Y cells into cells with morphological and biochemical characteristics of mature neurons. J Alzheimers Dis. 2010

24. Encinas M, Iglesias M, Liu Y, Wang H, Muhaisen A, Ceña V, Gallego C, Comella JX. Sequential treatment of SH-SY5Y cells with retinoic acid and brain-derived neurotrophic factor gives rise to fully differentiated, neurotrophic factor-dependent, human neuron-like cells. J Neurochem. 2000

25. Krishna S, Prajapati B, Seth P, Sinha S. LncRNA BASP1-AS1 is a positive regulator of stemness and pluripotency in human SH-SY5Y neuroblastoma cells. Biochem Biophys Res Commun. 2024

26. Koshy AM, Mendoza-Parra MA. Retinoids: Mechanisms of Action in Neuronal Cell Fate Acquisition. Life (Basel). 2023

27. Katsetos CD, Herman MM, Mörk SJ. Class III beta-tubulin in human development and cancer. Cell Motil Cytoskeleton. 2003

28. Witte H, Bradke F. The role of the cytoskeleton during neuronal polarization. Curr Opin Neurobiol. 2008

29. Zage PE, Nolo R, Fang W, Stewart J, Garcia-Manero G, Zweidler-McKay PA. Notch pathway activation induces neuroblastoma tumor cell growth arrest. Pediatr Blood Cancer. 2012

30. Szemes M, Melegh Z, Bellamy J, Greenhough A, Kollareddy M, Catchpoole D, Malik K. A Wnt-BMP4 Signaling Axis Induces MSX and NOTCH Proteins and Promotes Growth Suppression and Differentiation in Neuroblastoma. Cells. 2020

31. Sobecki M, Mrouj K, Camasses A, Parisis N, Nicolas E, Llères D, Gerbe F, Prieto S, Krasinska L, David A, Eguren M, Birling MC, Urbach S, Hem S, Déjardin J, Malumbres M, Jay P, Dulic V, Lafontaine DLj, Feil R, Fisher D. The cell proliferation antigen Ki-67 organises heterochromatin. Elife. 2016

32. Sun X, Kaufman PD. Ki-67: more than a proliferation marker. Chromosoma. 2018

